# An Optimized Approach for Annotation of Large Eukaryotic Genomic Sequences using Genetic Algorithm

**DOI:** 10.1101/083238

**Authors:** Biswanath Chowdhury, Arnav Garai, Gautam Garai

## Abstract

Detection of important functional and/or structural elements and identifying their positions in a large eukaryotic genome is an active research area. Gene is an important functional and structural unit of DNA. The computation of gene prediction is essential for detailed genome annotation. In this paper, we propose a new gene prediction technique based on Genetic Algorithm (GA) for determining the optimal positions of exons of a gene in a chromosome or genome. The correct identification of the coding and non-coding regions are difficult and computationally demanding. The proposed genetic-based method, named Gene Prediction with Genetic Algorithm (GPGA), reduces this problem by searching only one exon at a time instead of all exons along with its introns. The advantage of this representation is that it can break the entire gene-finding problem into a number of smaller subspaces and thereby reducing the computational complexity. We tested the performance of the GPGA with some benchmark datasets and compared the results with the well-known and relevant techniques. The comparison shows the better or comparable performance of the proposed method (GPGA). We also used GPGA for annotating the human chromosome 21 (HS21) using cross species comparison with the mouse orthologs.

## INTRODUCTION

Biological sequences are primarily useful computational data in molecular biology. Sequences represent symbolic descriptions of the biological macromolecules like DNA, RNA, and Proteins. A sequence can provide a vital insight into the biological, functional, and/or structural data about a molecule encoded in it. Therefore, the molecular information can be easily deciphered by analyzing several biological sequences. Over the past decade a major boost in sequencing, especially after the advent of *next-generation sequencing (NGS)* technologies (Liu et al. 2012) led to an enormous amount of nucleotide sequence data. Hence, the amount of raw, unannotated nucleotide sequence data in the databases is expanding exponentially. Understanding the functional significance of these data is the primary problem in comparative genomics. The use of computational approaches to accurately predict the functional and structural information of these DNA sequence data is an urgent requirement. Gene is the most important functional and structural unit of DNA. Hence, the computation of gene prediction is an essential part of the detailed genome annotation.

In an organism, DNA works as a medium to transfer information from one generation to another. A gene is a distinct stretch of DNA that determines amino acid residues of a protein or polypeptide, which are responsible for the biological functions in an organism. A gene undergoes transcription and translation process along with splicing to form a functional molecule. Most of the non-coding parts of DNA are spliced out during the process of transcription to form mature RNA from DNA. Three consecutive nucleotides or codon of a gene represents a single amino acid of a protein. A complete gene length is, therefore, always the multiplier of three. The prokaryotic gene structure contains long stretches of coding regions where intermediate non-coding regions are absent. On the other hand, the eukaryotic gene structure is more complex as it breaks into several coding regions or exons separated by long stretches of non-coding regions i.e. introns. Introns are spliced out from the transcribed RNA. Further, the coding region comprises only 2 – 3 % of the entire genomic sequence that adds a second level of complexity to identify a gene in eukaryotes. As a consequence, the gene prediction in a eukaryotic genome is more challenging. Researchers are attempting to get an efficient prediction tool that can accurately predict the location of genes in an unknown genomic sequence. However, the research towards the development of eukaryotic gene prediction algorithms is still yet to reach satisfactory results.

Computational gene finders have been able to predict genes precisely for single gene sequences, but for multi gene sequences, it lowers the accuracy with the increase of complexity that results false predictions. Research is still in progress to predict all exons correctly and subsequently to reduce the false predictions substantially (Guigó et al. 2000). Some of the well-known techniques available for gene prediction are GenScan (Burge and Karlin 1997), Genie (Reese et al. 2000), FGENESH (Salamov and Solovyev 2000), GeneId (Parra et al. 2000), GeneParser (Snyder and Stormo 1995), GRAIL II (Xu et al. 1994) and HMMgene (Krogh A 1997). Most of these prediction tools are based on classical approaches of gene identification such as Hidden Markov Models (HMM) (Kulp et al. 1996), Dynamic Programing (DP) (Jiang and Jacob 1998; Mott R 1997), and Bayesian methods (Pavlovic et al. 2002). Other than these, some of the approaches based on neural network (Zhou et al. 2008), wavelet transform (Abbasi et al. 2011), Genetic Algorithm (GA) (Sree and Babu 2013; Hwang et al. 2013) have been applied for accurate detection of a gene.

The ab-initio based method predicts the genes directly from the genomic sequences relying on two significant features like gene signals and gene content. Several ab-intio programs have been extensively used in genome annotation, such as GENSCAN, FGENESH, and GeneID. However, the ab-intio based approaches normally predict a higher rate of false positive results while annotating large multi genomic sequences (Dunham et al. 1999). In particular, ab-initio gene identifiers determine the intergenic splice sites poorly in the prediction process. Conversely, a homology search on the databases of already established and experimentally verified coding sequences have performed well and proved a good approach in decoding the structure of the genes having known homologs. At present, a large number of known protein coding genes, cDNA, proteins, and ESTs are available in the databases. Therefore, sequence similarity based gene prediction methods are becoming increasingly useful in finding the putative genes in genomic sequences and thereby provide an evolutionary relationship between the raw genomic data and known cDNA, proteins or gene databases.

To date, many different techniques are available for solving gene location detection and its structure prediction in the large eukaryotic genome. Acencio and Lemke (2009) introduced a decision tree-based classifier and trained that with different attributes like network topological features, cellular compartments, and biological processes to generate various predictors for finding essential genes in *S. cerevisiae.* EVidenceModeler (EVM) (Haas et al. 2008) was presented as a tool for automated eukaryotic gene structure annotation that computed weighted consensus gene structures based on both type and abundance of available evidence. SCGPred (Li et al. 2008) was another score based gene-finding program that combines multiple sources of evidence. Logeswaran et al. (2006) had developed a WAM-CpG algorithm based on the weight array method (WAM) and CpG islands. Nasiri et al. (2011) analyzed the performance of different ab-initio gene finders on orthologous genes of human and mouse. Genome Annotation based on Species Similarity (GASS) (Wang et al. 2015) is a shortest path model with DP based tool. It annotated a eukaryotic genome by aligning the exon sequences of the annotated similar species.

DNA numerical representation is another approach that was utilized in several algorithms where residues were converted to numerical values. Akhtar et al. (2008) had presented DNA symbolic-to-numeric representations and compared it with the existing techniques in terms of accuracy for both the gene and the exon prediction. Abbasi et al. (2011) showed a significant improvement in accuracy of exonic region identification using a signal-processing algorithm that was based on Discrete Wavelet Transform (DWT) and cross-correlation method. Saberkari et al. (2013) predicted the locations of exons in DNA strand using a Variable Length Window approach based on z-curve. It was a 3-D curve used to illustrate DNA sequences and to present a complete description of DNAs’ biological behavior. A Digital Signal Processing (DSP) based method was used by Inbamalar and Sivakumar (2015) to detect the protein coding regions where the DNA sequences were converted into numeric sequences using Electron Ion Interaction Potential (EIIP).

Evolutionary algorithms like GA based techniques have also been used in solving the gene prediction problem. Perez-Rodriguez and Garcia-Pedrajas (2011) developed a method based on genetic algorithm for gene prediction. Another GA approach (GA_PAUC) was used by Hwang et al. (2013) to maximize the partial Area Under the Curve (AUC). This technique used features of sequence information, protein-protein interaction network topology, and gene expression profiles to maximize the AUC of Receiver Operating Characteristic (ROC) plot. Amouda et al. (2010) proposed a web based tool Intron Multiple Aligner by Genetic Algorithm (iMAGA) for aligning the intron sequences to find their pattern. A model called feature-based weighted Naive Bayes model (FWM) which was based on Naïve Bayes classifiers, logistic regression, and genetic algorithm, was developed by J. Cheng et al. (Cheng et al. 2013).

In this paper, we propose a GA based optimized gene prediction method named as *Gene Prediction with Genetic Algorithm (GPGA).* It is used in the analysis of large, unknown eukaryotic genomic sequences by mapping with known genes. The advantage of this algorithm is that it can be utilized as a tool for identifying a gene optimally in a large genomic sequence. The GPGA is a novel evolutionary process with significant accuracy that can be utilized in mapping of a genome with genes present in several well-known repositories like *Ensemble* (http://www.ensembl.org), *UCSC* (http://www.genome.ucsc.edu) browser and others. We observe the performance of GPGA, which is very promising in terms of sensitivity and specificity on different benchmark datasets.

## RESULTS AND DISCUSSION

The performance of the GPGA was tested on two benchmark datasets, HMR195 (Rogic et al. 2001), and SAG (Guigó et al. 2000). In the experiment, we statistically evaluated the sensitivity and specificity of the proposed method at exon level and also compared the results with other well-known and relevant techniques. Further, we annotated human chromosome 21 with GPGA for large-scale evaluation.

The proposed algorithm has been written in C and implemented on an IBM Power 6 system with 8 GB RAM per core.

### Test datasets

To test the performance of GPGA, we considered two benchmark datasets of different categories having well-annotated genomic sequences. One dataset (HMR195) comprises the real genomic sequences where each sequence contains only one gene. Whereas, another dataset (SAG) consists of a set of annotated gene sequences which are arbitrarily placed in the background of random intergenic DNAs and each sequence contains more than one gene. The datasets were taken from the *GeneBench suite* (http://www.imtech.res.in/raghava/genebench). Brief descriptions of these datasets are given below.

*HMR195 dataset* comprises real genomic sequences of *H. sapiens, M.musculus, and R. norvegicus* in the sequence ratio of 103:82:10. Each sequence contains exactly one gene. The mean length of total 195 sequences is 7096 bp. The total number of single exon genes and multi-exon genes are 43, and 152, respectively. The number of exons in the dataset is 948 in total.

*SAG dataset* is the second one tested in the experiment. It consists of a semi-artificial set of genomic sequences with 43 simulated intergenic sequences. The dataset was developed by arbitrarily embedding a typical set of annotated 178 real human genomic sequences (h178) in those 43 sequences. Each of h178 sequences codes for a single complete gene. The SAG sequences have an average length of 177160 bp with 4.1 genes per sequence. The dataset contains total 900 exons.

### Data preprocessing (selection of homolog sets)

The GPGA is a GA-based homology technique, which determines the presence of a gene by identifying the position(s) of exons in a large unannotated eukaryotic DNA sequence. In the experiment, we compared the positions of exons found by the GPGA in the corresponding genomic sequence with the actual positions mentioned in the annotation file provided with the test datasets. For such test, we generated a customize dataset of homologous genes consulting both the test datasets (HMR195 and SAG).

We considered top three species, namely, human, mouse, and rat in searching the homologs for the test datasets since they are phylogenetically very close and the test datasets also consist of the genomic sequences of those three species. To construct a dataset of homologous genes for the sequences of test datasets, we used Blast Like Alignment Tool (BLAT) (Kent WJ 2002) of UCSC genome browser using the default nucleotide alignment parameters. We considered BLAT as the target database of BLAT is not a set of sequences, but instead an index derived from the assembly of the entire genome. First, we extracted the genes from the genomic sequences of both test datasets based on the positions of exons, mentioned in their respective annotation files and then make each of them as a query to the BLAT run. For HMR dataset, all 195 genes and for SAG dataset, 178 genes were searched against human, mouse, and rat genome separately using their latest assembly (Human: hg38; mouse: mm10; and rat: rn6). For human origin of the test genomic sequence, we searched homologous sequences in mouse and rat, for mouse origin, we searched homologs in human and rat, and for rat origin we searched in human and mouse. We included the highest scored homologous sequence from the BLAT search. Thus, we chose two homologous sequences of the other two species for each query gene and the sequence of the query itself. The execution of GPGA had not been performed directly with the extracted exons from the genomic sequences of test datasets based on the mentioned positions in the annotation file. Because position comparison of GPGA by extracting exons based on the actual position with the given annotated position reduces the real genomic level complexity. In BLAT run, although we had always considered the top homologs, some of them are of poor quality in terms of similarity. It is also noted that some of the homologous sequences did not contain the same number of exons of the query and/or precise exon boundary presumably because the BLAT results contain newly assembled genome with respect to our benchmark datasets. However, we included those sequences to our new homolog sets to increase the noise in gene data and to test the efficiency of GPGA by excluding them as falsely verified sequences. Duplicate inclusion of a same homologous sequence for different query (gene) was eliminated from the sets. Thus, we prepared two homolog sets, one for HMR dataset and another for SAG dataset and finally we combined them to prepare one customized homolog set. The process flow for generating the final homolog dataset is shown diagrammatically in Figure 1.

**Figure 1:**
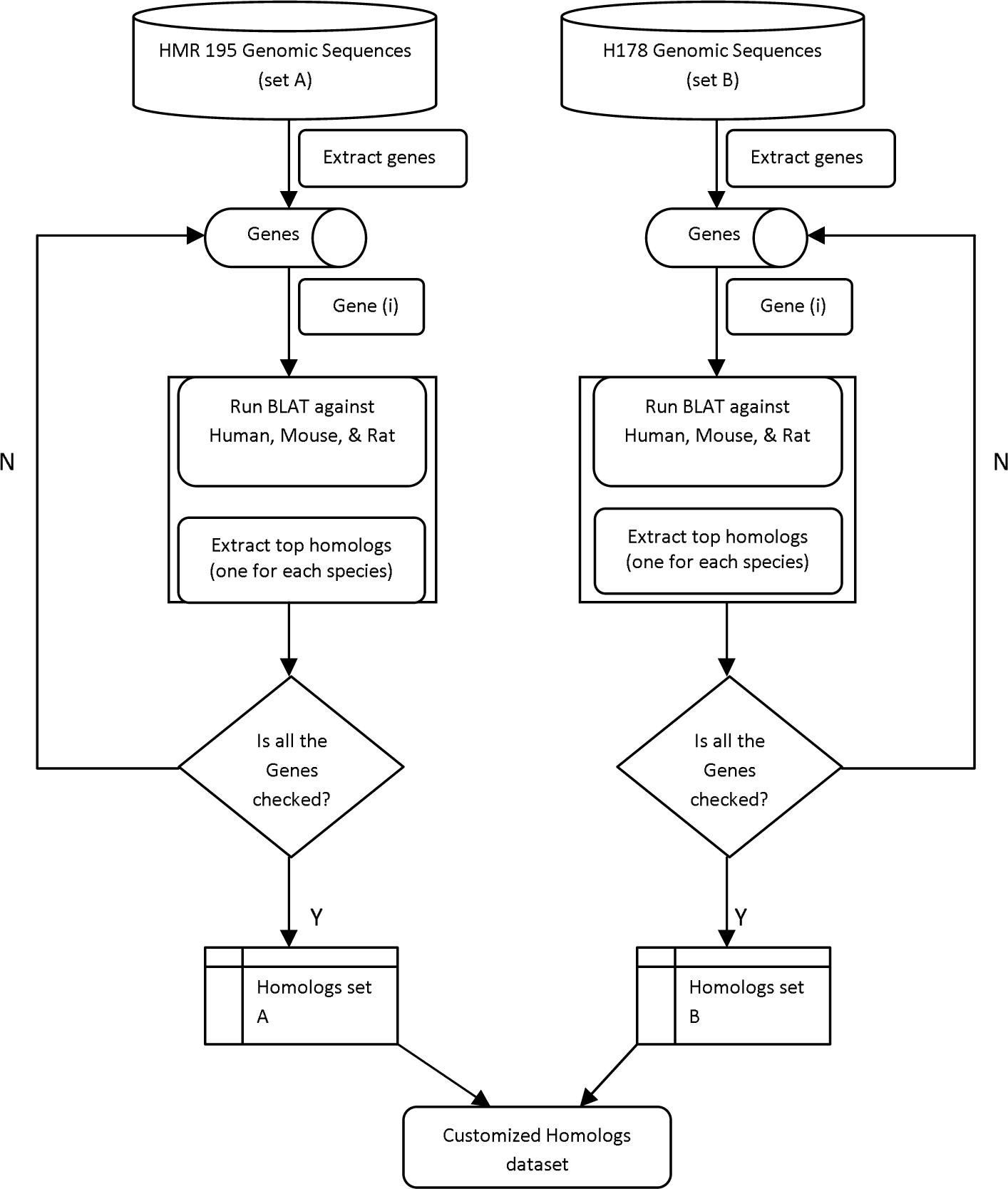
The flow chart representing the process of homologs dataset construction

The proposed method extracted each of the exons separately from the homolog sets and searched for the presence of them to 195 genomic sequences of HMR and 43 sequences of SAG considering both plus (Watson) and minus (Crick) strands.

### Performance assessment

The predicted exon positions by GPGA were compared with the actual exon positions present in their corresponding annotation file. In the post-processing of matched result, we had considered a filtering criterion to identify a gene as true prediction where minimum 60 percent similarity at the gene level along with at least 60 bp sequence length was found. We performed statistical analysis of the experimental results to determine the performance accuracy of GPGA (see Methods for details). The results were also compared with other well-known and relevant annotation tools.

For both HMR and SAG datasets, we measured *ESn* (sensitivity at the exon level), *ME* (missed exon), *ESp*, (specificity at the exon level), and *WE* (wrong exon) using human, mouse, and rat homolog sets separately. The average value of each parameter was calculated separately for human, mouse, and rat homolog sets and was considered as the final measurement (see Supplemental_file_1.pdf: Statistical analysis and Table S1). The statistical measure was not considered when GPGA did not find any such homologous exons of a gene in a sequence. Figures 2 and 3 (Tables S2 and S3 in Supplemental_file_1.pdf) show the comparison of the GPGA results with other well-known gene prediction tools on HMR and SAG datasets, respectively.

**Figure 2:**
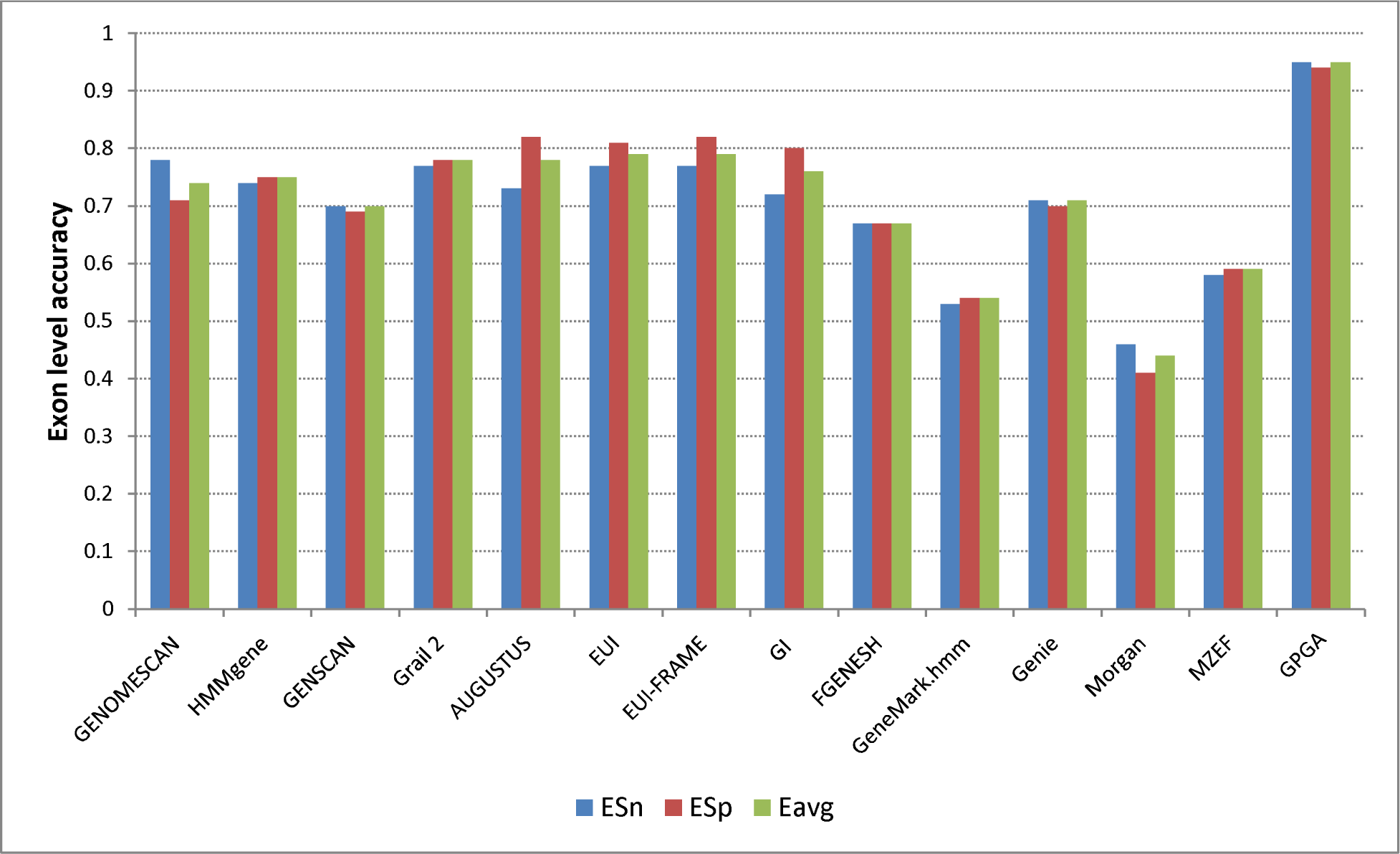
The exon level accuracy comparison of GPGA with other gene prediction tools on HMR dataset

**Figure 3:**
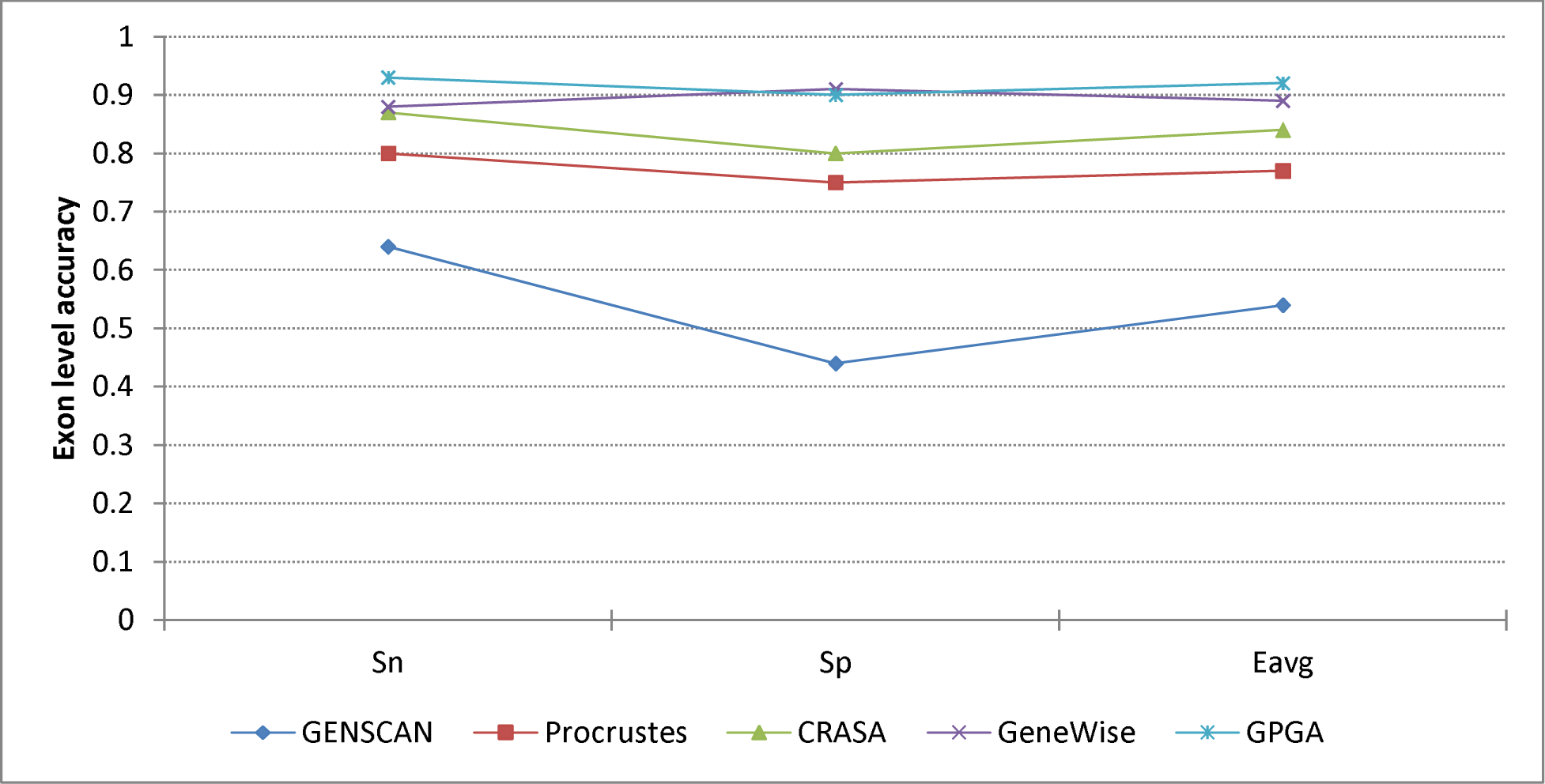
The exon level accuracy comparison of GPGA with other gene prediction tools on SAG dataset

Figure 2 depicts the high accuracy of the GPGA performance on HMR dataset in terms of *ESn, ESp*, and *Eavg*, and it consistently was above 90% for both *ESn* and *ESp.* The values of *ESn, ESp*, and *Eavg* of GPGA are 0.95, 0.94, and 0.95, respectively. From the comparison, it is noticed that GPGA outperformed the other tools significantly.

Figure 3 also illustrates a better performance of GPGA compared to other annotation tools in terms of sensitivity and specificity at exon level. It was noted that GPGA performed similar to GeneWise. However, the (93%) of GPGA is better than *ESn* (88%) of GeneWise and the *ESp* of GPGA and GeneWise are 90% and 91%, respectively. However, the overall consistency of GPGA (*Eavg* = 0.915) is higher than GeneWise (*Eavg* = 0.89).

In addition to *ESn* and *ESp*, for measuring accuracy, *ME* (the proportion of missing exons and actual exons) and *WE* (the proportion of predicted wrong exons and actual predicted exons) were also included in the evaluation process for the superiority of the tools. Here, the GPGA also performed well. The accuracy measurement parameters are presented in Supplemental_file_2.xls (see Table S4 and S5). Furthermore, even when a good similarity is found, the limits of predicted exon positions were not always very precise. Small exons were also missed by GPGA because of the presence of other alternative regions. We showed the performance of GPGA in terms of wrong exon prediction for both datasets in Figure 4.

**Figure 4:**
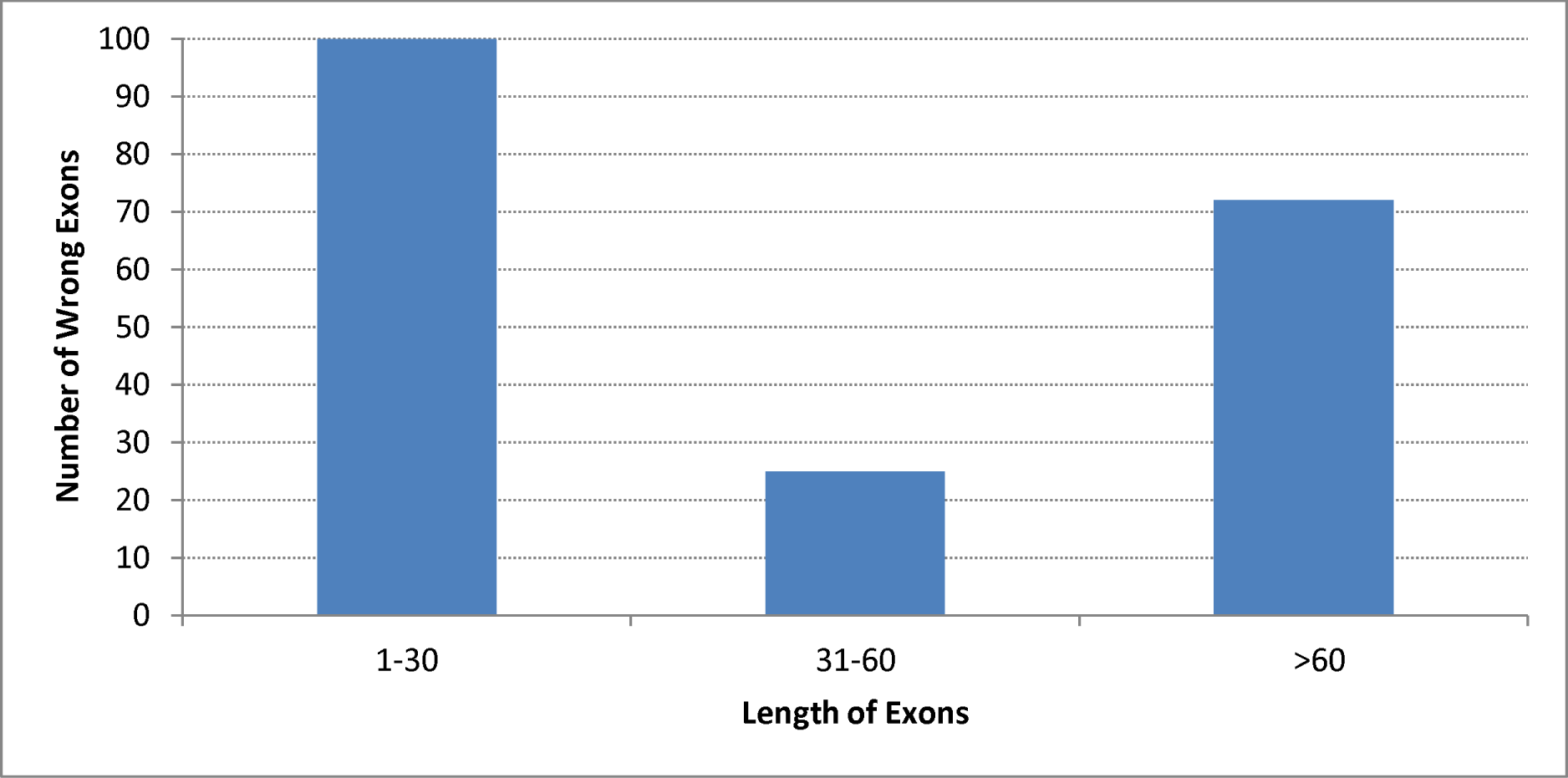
Total number of wrong exon prediction by GPGA at different range of exon length

From Figure 4, it is observed that the number of wrong exon prediction is maximum for the smallest length range exons (≤30). This is because for a small length exon, the chance of getting alternative regions, other than the actual position is higher.

### Annotation of human chromosome 21

We had also performed annotation of human chromosome 21 (HS21) to observe the performance of GPGA at the chromosome level. We have selected HS21, as it is the smallest human autosome that wraps around 1-1.5% of the human genome and its structure and gene content have also been intensively studied. Therefore, it is considered as an excellent dataset to validate any gene prediction method. Number of gene identification on each human chromosome using various approaches is now an active research area. One of the most challenging works among them is the cross-species comparison of human genome. The comparison of human genome with other related species helps to identify the related genes since the functionally related sequences tend to be actively conserved through evolution. We have chosen the phylogenetically related species like mouse genome for cross-species comparison with the human genome. HS21 shows conserved syntenies to mouse chromosomes 16, 17, and 10 (MM-10, MM16, and MM17) (http://www.ncbi.nlm.nih.gov/projects/homology/maps/human/chr21). Hence, we selected sequences from MM10, MM16, and MM17. We then mapped mouse DNA sequences onto HS21 in order to identify regions that have been conserved.

#### Data Pre-processing (selection of target and reference sequences)

The target sequence for this analysis is HS21, and the reference sequences are MM10, MM16, and MM17. The entire HS21 sequence of ~47 mb (GRCh38.p4) along with its seven alternate loci (ALT_REF_LOCI_1) was obtained from the NCBI (http://www.ncbi.nlm.nih.gov/assembly/GCF_000001405.30). Alternate loci are multiple representations for regions that are too complex to be represented by a single path (https://www.ncbi.nlm.nih.gov/projects/genome/assembly/grc/human/). The main objective was to map the target sequence, HS21-specific genes with their reference mouse orthologs. We analyzed non-repetitive parts of the HS21 sequence by aligning with well-annotated mouse CoDing Sequences (CDSs) of MM10, MM16, and MM17. The reference coding sequences were obtained from the Gencode assembly using UCSC browser. GENCODE Comprehensive set is richer in alternative splicing, novel CDSs, novel exons and has higher genomic coverage than RefSeq while the GENCODE Basic set is very similar to RefSeq. Thus, we selected comprehensive Gencode VM4 published in Aug, 2014.

Due to limited computing resources, we initially divided the target sequence into multiple (total 26 numbers) divisions. Each of them consists of 16-lac bp of HS21 except the last one. Each of the divisions was run against total comprehensive sets of MM10, MM16, and MM17.

#### Results of Annotation

In the experiment, we analyzed the results defining the stringency based on the lengths and similarities of the conserve sequences. We accordingly categorized the sequence length into 50, 100, and 150 bp. For each sequence length, we considered four types of percentage similarity, i.e., 60, 70, 80, and 90. For each category of length along with its similarity, we found a large number of conserved blocks. A gene is considered to be conserved between human and mouse if all the exons of a gene were matched significantly. For e.g., for the threshold criteria of 100 bp with 60% similarity, a gene is considered conserved if there is 60% similarity for all mouse exons. Even if a single exon satisfied the threshold criteria; we considered it as a conserved block. However, that exon might be or might not be a part of a gene. We found a large number of such blocks. This presumes the presence of a large number of potentially functional, non-genic conserve regulatory and/or structural blocks. Figures 5a and 5b, respectively, show the ungapped conserved blocks distribution and the total number of genes for different sequence lengths and similarity categories. Out of different categories, we had chosen the stringency of 100 bp with 70% identity (represented as 100-70) as our final criteria to find ungapped conserved blocks and genes. Below this threshold value, we identified a large number of conserved blocks and genes. However, a large number of exons of a gene did not follow GT-AG splicing rule. We also got a significant number of blocks and genes over 150 bp stringency. However, we considered medium length of 100 bp as our final value to balance the sensitivity as well as specificity (details are provided in Supplemental_file_3.pdf: Table S6). Following the stringency of 100-70, (Table 1) yielded 2136 conserve blocks and 361 homologous genes for HS21. Those 361 genes contain total 3150 exons. Out of them, 2185 exons are with canonical ‘GT-AG’ splicing junctions and 604 with non-canonical ‘GT-AG’ junctions. It was also observed that out of the 361 genes, 63 genes are overlapping genes (where both ends were not mapped by the mouse orthologs) and 149 are partial genes (having only one end matched). The GC percentage was 51.68, which was a significant one. Considering pseudogenes based on retroposen, and the genes having premature stop codon we found 41 genes. The distribution of blocks and genes along the length of HS 21 is shown in Figure 6 (see Supplemental_file_3.pdf: Table S9). From Figure 6, it was noted that the regions of conserved blocks and the locations of genes were close to each other and they were distributed more at the distal part (gene-rich region) of HS21. When the sequences were compared with 80% identity over 100 bp (100-80), 1607 conserve blocks and 194 genes were detected.

**Figure 5:**
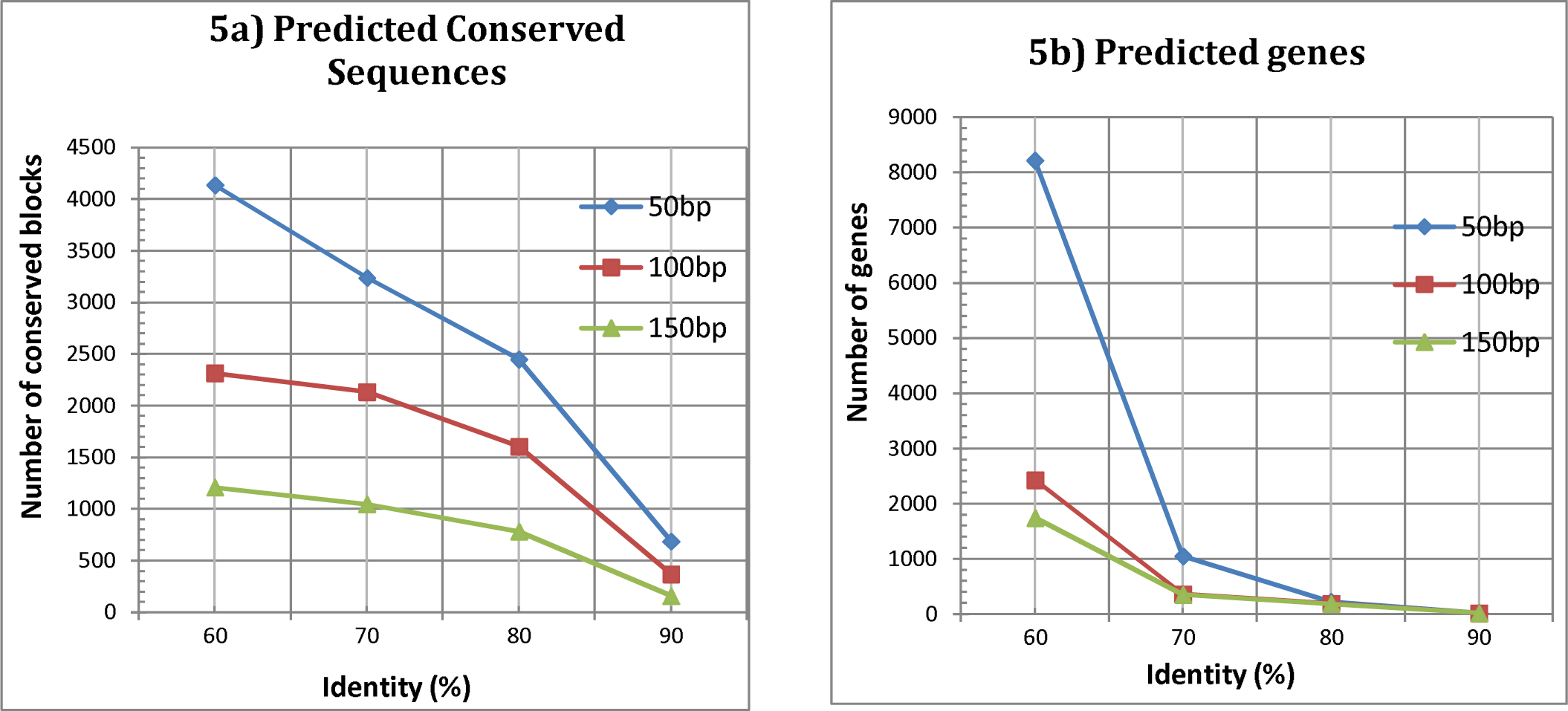
Results of Conservation identified by GPGA based on different threshold criteria a) Number of ungapped conserved blocks b) Number of genes.

**Table 1.** Results of GPGA for Human Chromosome 21.

| Stringency at 100 bp length with 70% similarity | HS21 |
| --- | --- |
| 1. Total number of conserve blocks | 2136 |
| 2. Total number of genes (including partial, overlapping, and retroposen) | 361 |
| 2.1. Total number of exons in all genes | 3150 |
| 2.2. Number of GT-AG junctions | 2185 |
| 2.3. Number of non GT-AG junctions | 604 |
| 2.4. Total number of residues comprising all the genes | 412168 |
| 2.5. Total number of partial genes that have 5' end matched | 77 |
| 2.6. Total number of partial genes that have 3' end matched | 72 |
| 2.7. Total number of overlapping genes | 63 |
| 2.8. Total number of retroposen (may include partial or overlap genes) | 41 |
| 2.9. GC percentage | 51.68 |

**Figure 6:**
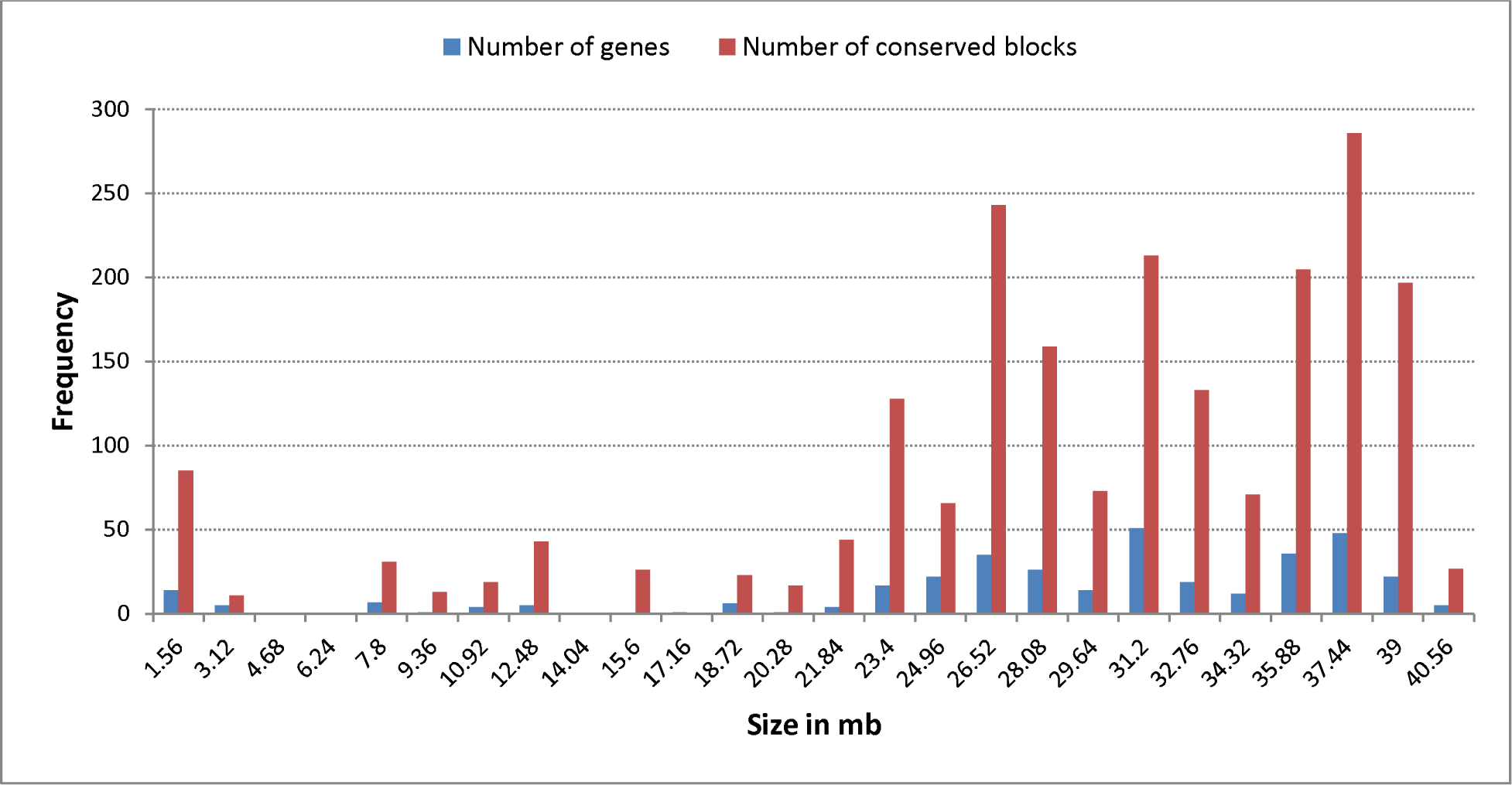
Distribution of conserved blocks and genes all along the human chromosome 21

For 100-70 level, we also provided the base substitution data in Supplemental_file_3.pdf (seeTable S7, S8, and Figure S1) showing the highest rate of transitions (substitution between two purines and between two pyrimidines) than transversion (substitution between one purine and one pyrimidine) and the higher rate of substitution at third codon position (Wobble position) than first and second.

To compare the GPGA result with others, we considered only those genes that have either unique start or end positions. We excluded alternate transcripts having the same start and end positions. Out of the 361 genes predicted by GPGA for HS21, we found 283 genes share unique start and/or end positions. Table 2 shows the comparative results of GPGA with Refseq and Gencode assembly. Refseq found 411 genes, of which 271 contain unique start and/or end positions. Out of 271 genes, 158 genes were partially (either of the both ends of a gene) predicted by the GPGA. The partial prediction is because, GPGA performed ungapped mapping for finding the conserved genes, whereas, the results of other methods showed alignment including gaps. Gencode basic assembly got 309 unique genes, of which 162 genes were partially predicted by the GPGA. On the other hand, Gencode comprehensive set got more genes than Gencode basic as it contained more novel exons. It predicted 509 unique genes, out of which 174 genes were partially predicted by GPGA. Table 3 contains the comparative results of GPGA along with other gene prediction tools. From the table, it is noticed that the GPGA predicted more genes than other gene prediction tools.

**Table 2.** Comparative results are showing the different assembly along with matching genes with GPGA prediction.

| Assembly | Total genes | Total genes crossed 100-70 threshold level | Total genes with either unique start/end position | Number of genes matched with GPGA prediction |
| --- | --- | --- | --- | --- |
| Refseq | 479 | 411 | 271 | 158 |
| All_GencodeV22_Basic | 609 | 515 | 309 | 162 |
| All_GencodeV22_Comprehensive | 896 | 739 | 509 | 174 |
| All_GencodeV23_Basic | 607 | 515 | 309 | 162 |
| All_GencodeV23_Comprehensive | 896 | 740 | 511 | 174 |
| GencodeV20_basic (Ensemble76) | 602 | 502 | 302 | 163 |
| GencodeV20_comprehensive (Ensemble76) | 883 | 721 | 499 | 176 |
| GPGA Genes | 361 | 361 | 283 | . |

**Table 3.** Comparative results are showing the different annotation tools along with matching genes with GPGA prediction.

| Gene prediction tools | Total genes | Total genes crossed 100-70 threshold level | Total genes with either unique start/end position | Number of genes matched with GPGA prediction |
| --- | --- | --- | --- | --- |
| CCDS | 339 | 287 | 238 | 149 |
| AUGUSTUS | 248 | 181 | 126 | 82 |
| GeneID | 271 | 122 | 122 | 85 |
| GENSCAN | 420 | 77 | 77 | 43 |
| SGP Genes | 271 | 203 | 203 | 123 |
| GPGA Genes | 361 | 361 | 283 | . |

The results proved the performance superiority of GPGA compared with other well-known ab-initio or homology based approaches. Nevertheless, due to limited computational resources, we split the entire HS21 into several segments. This approach increased the computation time.

## CONCLUSION

The proposed approach (GPGA) is an integer based evolutionary process which simplifies the gene prediction technique. The GPGA was tested with two well-known benchmark datasets HMR195 and SAG to evaluate the performance in terms of sensitivity and specificity at the exon level. However, one of the datasets HMR195 consists of real genomic sequences and the other one SAG contains semi-artificial set of genomic sequences. Such choice of datasets helps to truly measure the performance of an approach in a noisy environment. Finally, it was noted that the GPGA truly predicted the gene better than other well-known approaches and its accuracy is more than 90%.

The limitation of GPGA is that it often fails to predict the correct position of a short length exon since same sequence is frequently repeated in a genomic sequence. Another shortfall of GPGA is that it can perform well on an unannotated raw sequence, only when there is a well coverage of annotated information of orthologous genes.

In future research work, we want to introduce further the information of content sensors and signal sensors like GC-content value, TATA box, promoters and other compositional parameters along with the sequence homology to improve the sensitivity of the GPGA. We also wish to perform parallel computing for large-scale annotation without splitting the query length. In addition, we would like to observe the performance of the GPGA after introducing gaps in it.

## METHODS

### Genetic algorithm

One of the most commonly used evolutionary techniques for optimization is GA, which is stochastic in nature. It iteratively executes a set of individuals called a population. Each individual is referred to as a chromosome that encodes a possible solution to the given problem. Each solution is assigned a problem specific fitness score. After every iteration (generation), the fittest individuals are carried on to the next generation, and this process continues until a termination criterion is fulfilled. The three genetic operators – selection, crossover, and mutation help to modify a population in each generation. The conventional GA normally represents a chromosome by a binary string. Binary representation, however, can be problematic for solving some problems as it is sometime difficult to encode a real problem with binary window. Another problem in binary coding is the increased length of the string for representing a large and complex optimization problem, which increases the computational complexity and the memory space. So, the problem specific GAs have been necessary to develop with other types of representations in mind, apart from binary notation.

One of the most used GAs is the Real coded GA (RGA), whose significance is justified in several theoretical studies (Goldberg DE 1991; Radcliffe NJ 1991). In RGA, chromosomes are represented by the real (floating) numbers instead of binary numbers. Moreover, the researchers have suggested several modifications to the GA operators other than conventional one point crossover, two point crossover, bitwise flip mutation (Goldberg DE 1991). Among them, few examples of the crossover operation in GA are flat crossover (Radcliffe NJ 1991), arithmetical crossover (Michalewicz Z 1996), BLX-α (Eshelman and Schaffer 1993), Laplace crossover (Deep and Thakur 2007a), Simulated Binary Crossover (SBX) (Deb and Agrawal 1995), aligned block crossover (Garai and Chowdhury 2015). Similarly, the modifications in mutation operator like polynomial mutation (Deb and Agrawal 1999), uniform and non-uniform mutation (Michalewicz et al. 1994), power mutation (Deep and Thakur 2007b), controlled mutation (Garai and Chowdhury 2015) have been used to improve the GA process depending on the application problem.

Here, we have modified the conventional GA with the integer coding. The changes in crossover and mutation have also been performed for solving the problem efficiently. Such modification improves the performance of the proposed GPGA.

### Gene Prediction with Genetic Algorithm

The objective of the proposed method (GPGA) is to map a well-annotated known sequence onto an unknown large genomic sequence. The mapping determines the homologous relationship between the known sequence (with the known genes) and the unknown genomic sequence by identifying the homologous gene(s) in the unknown sequence. Eukaryotic gene contains long stretches of introns that intervenes the coding parts or exons. CDS is composed of exons, which actually the translated portion of a gene. CDSs are the important parts of genes and are structurally more conserved in homologous sequences. However, to find the small and discrete portions of CDS in a large genomic sequence is an exhaustive search procedure and requires a significant amount of computational time and memory space. We have incorporated an integer based GA (IGA) approach in GPGA to overcome such problems. Gene representation by GPGA

In the proposed method, the individuals of the GA population are represented by integer values. These values signify different possible positions of an exon in a large unknown genomic sequence. In GPGA, the searching process iteratively reaches the optimum position to define the actual position of the exon. As a result, instead of searching the entire gene (comprising a number of exons) in an unknown genome, the GPGA separately looks for each exon of the corresponding gene. Thus, the execution of GPGA is dependent on the number of exons present in a gene. The advantage of such representation is that it breaks up the search space of the gene-finding problem to a number of smaller subspaces and thereby reducing the computational complexity. It eventually reduces the possibility to be stuck up in a local optimum.

### Population Initialization

In the initialization step, an integer based initial population of size *N* is randomly generated within a lower and an upper limit. Each individual or a chromosome *P_i_, ∀ i ∈ {1, 2,…, N}* is an integer value that represents a probable starting location of an exon (*E*) in the query genomic sequence (*Q*). The lower and the upper limits define the lowest and the highest positions in *Q*, which are two constant starting positions of an exon. The lower limit (*l*) defines the starting position of *Q* i.e., 1. The upper limit (*u*) is the difference between the length of *Q* and the exon (*E*) length, i.e., if the length of *Q* is *q* and the length of *E* is *e*, the upper limit *u* is (*q − e*).

### Fitness Function

The fitness score of a chromosome represents the alignment score. The alignment finds the presence of a conserve region (exon) in the query sequence. In the score calculation, we have considered that an identical match gets +1, and a mismatch gets a 0. Thus, the score is computed by the following fitness function,

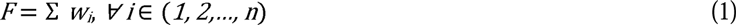

where, *w*_*i*_ defines a local alignment score and *n* is the total number of local alignments. *w*_*i*_>0, if any locally matched portion is found, otherwise, *w*_*i*_ = 0.

Therefore, the fitness value (*F*) of a chromosome denotes the summation of all local alignment scores. Now, let the chromosome be *P*_*1*_. The fitness score calculation of *P*_*1*_ is shown in Figure 7.

**Figure 7:**
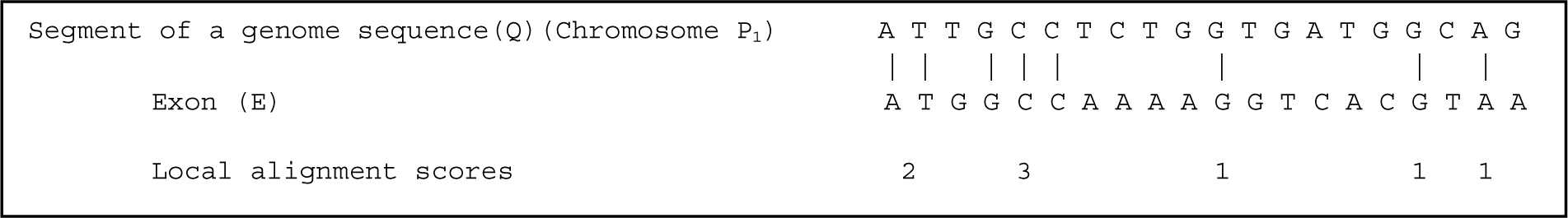
Fitness score calculation in GPGA

Figure 7 shows five local alignment scores for *P*_*1*_. According to the Equation 1, the fitness score of *P*_*1*_ will be (*P*_*1*_) = 2+3+1+1+1=8.

### Genetic operators

Three genetic operators namely, *selection, crossover*, and *mutation* play an important role towards the convergence of the problem. These operators also maintain a balance between the exploration and exploitation of the search space (Ortiz-Boyer et al. 2007).

#### Selection operator

In GPGA, we have considered tournament selection technique with tournament size 3 as a selection operator. In this approach three individuals are chosen randomly from the population pool *P_i_, ∀ i ∈ {1, 2,…, N}* and are entered into the tournament. Based on the fitness value, the fittest individual among three, say, *P*_*a*_ will be selected to take part in the crossover operation. This process is continued along with crossover and mutation until an entire new population *P’*_*j*_, *∀ j ∈ {1, 2,…, N}* is generated.

#### Crossover operator

In the GPGA, we have considered a modified crossover operation named as *Adaptive Position Prediction (APP)* crossover. APP crossover is a self-controlled-crossover operation that adaptively modifies *l* and *u* depending on the fitness score of parents. Let us consider two parents (say, *P*_*a*_ and *P*_*b*_) are randomly selected from the population pool. By this operation, two offsprings (say, *P’*_*a*_ and *P’*_*b*_) are generated from the selected parents. APP crossover depends on the fitness (alignment) scores of *P*_*a*_ and *P*_*b*_ However, the *maximum fitness score* of a parent will never exceed *e* (the length of the exon). If the score is *e*, then it is considered that the optimal exon (*E*) region is found and the exon is entirely overlapped. On the other hand, if the score is either significantly close to *e*, then the condition is called finding of the suboptimal exon region. Then a part of the exon (*E*) is overlapped and the APP crossover narrows down the range of limits *l* and *u* close to the parents to search for offsprings. The default cutoff score for a suboptimal exon region is selected as 50% of the *maximum fitness score*, i.e., *e*/2. Any fitness score less than *e*/2 is discarded. Now, let, the fitness score of *P*_*a*_ and *P*_*b*_ be *P*_*a*_^*obj*^ and *P*_*b*_^*obj*^, respectively. If *P*_*a*_^*obj*^ ≥ *e*/2 and *P*_*a*_^*obj*^*>P*_*b*_^*obj*^, then the offsprings *P’*_*a*_ and *P’*_*b*_ will be produced as follows.

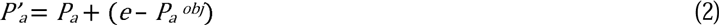

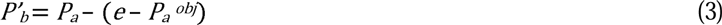

However, if *P*_*b*_^*obj*^ ≥ *e*/2 and *P*_*b*_^*obj*^>*P*_*a*_^*obj*^ then the offsprings *P’*_*a*_ and *P’*_*b*_ will be generated as below.

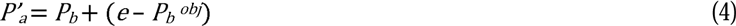

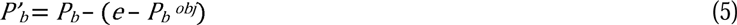

On the other hand, if *P*_*b (obj)*_ and *P*_*a (obj)*_ are less than *e*/2, then *P’*_*a*_ and *P’*_*b*_ are produced by choosing a random number θ between *P*_*a*_ and *P*_*b*_ as follows.

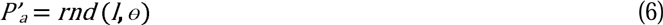

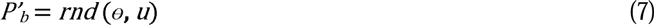

where, *rnd* is the function to generate random number. Thus, the crossover operation helps to predict the correct exon position by adaptively narrowing down the difference between *l* and *u.* This adaptive nature helps in fine-tuning of the operator for converging to the optimal position.

The APP crossover operation is represented algorithmically in the following way.

- Randomly select two parents *P*_*a*_ and *P*_*b*_ from a population pool having fitness scores *P*_*a*_^*obj*^ and *P*_*b*_^*obj*^, respectively.
- Two offsprings *P’*_*a*_ and *P’*_*b*_ are produced as follows: if ((*P*_*a*_^*obj*^ ≥ *e*/2 and *P*_*a*_^*obj*^>*P*_*b*_^*obj*^)) then do

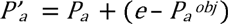

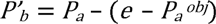

if (*P’*_*a*_>*u*) then do *P’*_*a*_ = *u* endif if (*P’*_*b*_<*l*) then do *P’*_*b*_ = *l* endif endif else if (*P*_*b*_^*obj*^ ≥ *e*/2) then do

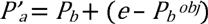

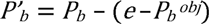

if (*P’*_*a*_>*u*) then do *P’*_*a*_ = *u* endif if (*P’*_*b*_<*l*) then do *P’*_*b*_ = *l* endif endif else Choose a random integer number θ between and *P*_*a*_ and *P*_*b*_

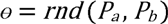 Then, generate two offsprings as follows

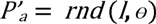

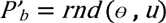 End

#### Mutation operator

The mutation operation is performed similar to the APP crossover. It is also named as Adaptive Position Prediction (APP) mutation. It mutates the offspring generated from the crossover operation to another possible offspring to maintain the diversity in the population for faster searching of the optimal position of the given exon (*E*). Let, the fitness score of an offspring *P’*_*a*_ be *P’*_*a*_^*obj*^. If *P’*_*a*_^*obj*^ ≥ *e*/2, then the modified new lower limit (*l*_*m*_) and new upper limit (*u*_m_) will be defined as follows.

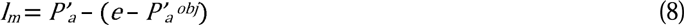

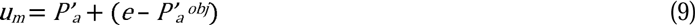

Thus, the modified offspring is generated as follows.

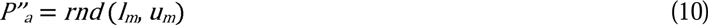

where, *rnd* is the random number generator. If *P’*_*a*_^*obj*^<*e*/2, then the offspring is modified by the APP mutation as follows.

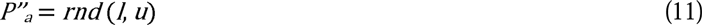

where, *rnd* is the random number generator.

The algorithmic steps of the APP mutation operation are given below.

- Select an offspring *P’*_*a*_.
- Mutate *P’*_*a*_ to *P”*_*a*_ as follows: if (*P’*_*a*_ *obj* ≥ *e*/*2*) then do

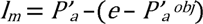

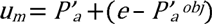

if (*l*_*m*_<*l*) then do *l*_*m*_ = *l* endif if (*u*_*m*_>*u*) then do *u*_*m*_ = *u* endif *P”*_*a*_ = rnd (*l*_*m*_, *u*_*m*_) Endif else do *P”*_*a*_ = (*l, u*) End

### Termination

The process is terminated when the maximum number of iterations *(generations), G*_*max*_ is reached. However, to reduce the computation time without compromising the accuracy level, another termination criterion based on the fitness score of the best individual is set. If the score of the best solution remains unchanged for 200 consecutive generations, then the process is stopped.

Now, the proposed GPGA is represented algorithmically in the following way.

1. Read the unknown genomic sequence (*Q*) and the reference exon sequence (known) (*E*) which is to be mapped.
2. Initialize the population size *N*, AAP crossover probability (*P*_*cross*_), AAP mutation probability (*P*_*mut*_) and *G* = 1
3. Generate an initial population *P*_*i*_, *∀ i ∈ {1, 2,…, N}* of *N* individuals(chromosomes). Where each chromosome represents a probable starting position of *E* in *Q*.
4. Evaluate the potential of each individual *P*_*i*_, *∀ i ∈ {1, 2,…, N}* in terms of fitness score based on the objective function *F* (discussed in Fitness Function).
5. Select individuals from the pool of *N* individuals using the tournament selection with *tournament size* 3 and pick up two best individuals *P*_*a*_ and *P*_*b*_ based on fitness value.
6. Perform the AAP crossover operation (discussed in *Crossover operator)* with *P*_*cross*_ between the selected individuals *P*_*a*_ and *P*_*b*_ and mutate them (discussed in *Mutation operator)* with mutation probability, *P*_*mut*_.
7. Each pair of individual (*P*_*a*_ and *P*_*b*_) generates two children *P’*_*a*_ and *P’*_*b*_.
8. Repeat steps 5 – 7 until a new pool of individuals *P’*_*i*_, *∀ i ∈ {1, 2,…, N}* is formed and *G = G + 1.*
9. Stop the process if the termination criterion is satisfied (discussed in Termination). Otherwise, go to step 4.

### GPGA parameters

In the proposed method, we considered the values of *N* = 200 and *G*_*max*_ = 3000. Since, the computational time increases with *G*_*max*_ value, we set the termination criterion based on the convergence of the best fitness score (see Termination). This approach always prevents the unwanted computation of GPGA upto *G*_*max*_. The optimum value of *N* was set to 200 as it produced the best results in the experiment. For GPGA, we allowed crossover and mutation operations to perform in every iteration or generation since it converges faster to an optimal solution. As a result, we set up *P*_*cross*_ = 1, and *P*_*mut*_ = 1. This eventually relieves the user to choose specific input values of *P*_*mut*_ and *P*_*cross*_. Thus, the user with less or no prior knowledge of the GA can run GPGA very easily without concerning about the optimal value of *p*_*cross*_ and *p*_*mut*_.

### Evaluation of prediction accuracy

Gene prediction accuracy of GPGA was computed at the level of exons. We followed the standard measures of sensitivity (*ESn*, and *ME*) and specificity (*ESp*, and *WE*) for evaluating the performance accuracy as described previously (Burset and Guigo 1996), and are formulated below.

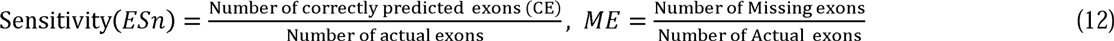

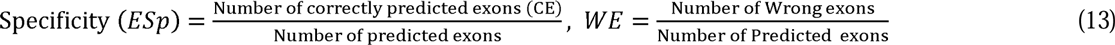

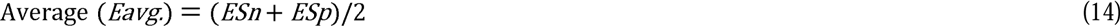

The predicted exon is regarded as correct only if its both sides’ boundaries are predicted correctly.

## ACKNOWLEDGMENTS

The authors would also like to thank Mr. R. Banerjee, S. Majumdar, and Mr. N. Sanpui for their helpful support in preparation of the manuscript.

## DISCLOSURE DECLARATION

### Competing interests

The authors declare that they have no competing interests.

### Ethics approval

No ethical approval was required for this study.

